# The allosteric activation of α7 nAChR by α-conotoxin MrIC is modified by mutations at the vestibular site

**DOI:** 10.1101/2021.04.14.439845

**Authors:** Alican Gulsevin, Roger L. Papke, Clare Stokes, Hue N. T. Tran, Ai-Hua Jin, Irina Vetter, Jens Meiler

## Abstract

α-conotoxins are 13-19 amino acid toxin peptides that bind various nicotinic acetylcholine receptor (nAChR) subtypes. α-conotoxin Mr1.7c (MrIC) is a 17 amino acid peptide that targets α7 nAChR. Although MrIC has no activating effect on α7 nAChR when applied by itself, it evokes a large response when co-applied with the type II positive allosteric modulator PNU-120596, which potentiates α7 nAChR response by recovering it from a desensitized state. Lack of standalone activity despite activation upon co-application with a positive allosteric modulator was previously observed for molecules that bind to an extracellular domain allosteric activation (AA) site at the vestibule of the receptor. We hypothesized that MrIC may activate α7 nAChR allosterically through this site. We ran voltage-clamp electrophysiology experiments and *in silico* peptide docking calculations to gather evidence in support of α7 nAChR activation by MrIC through the AA site. The experiments with the wild-type α7 nAChR supported an allosteric mode of action, which was confirmed by the increased MrIC + PNU-120596 responses of three α7 nAChR AA site mutants that were designed *in silico* to improve MrIC binding. Overall, our results shed light on allosteric activation of α7 nAChR by MrIC and suggest involvement of the AA site.

**Significance Statement:** α-conotoxin MrIC (MrIC) is an allosteric agonist of the α7 nicotinic acetylcholine receptor (nAChR). This mode of action is unique among α-conotoxins since these peptides typically act as orthosteric antagonists of nAChR. However, the mechanism of α7 nAChR activation by MrIC has been elusive so far. This work demonstrates that activation by MrIC is independent of the α7 nAChR orthosteric site and is related to a vestibular allosteric activation site at the extracellular domain of the receptor. Our experimental and computational studies identified the residues that play a role in allosteric activation and confirmed the utility of ensemble docking methods in understanding peptide – nAChR interactions, thus providing a basis for the design of peptides for allosteric modulation of nAChR.

## Introduction

### α7 nAChR structure and properties

The α7 nicotinic acetylcholine receptor (nAChR) is a homo-pentameric ligand-gated ion-channel belonging to the Cys-loop receptor family (1). α7 nAChR has unique properties among the nAChR including high calcium permeability when activated by conventional agonists (2), fast and concentration-dependent desensitization (3), and the presence of five putative orthosteric agonist binding sites (4). α7 nAChR have been targeted for the treatment of multiple diseases including depression (5, 6), schizophrenia (7, 8), Alzheimer’s disease (9), and for modulation of pain (10, 11). Most ligands targeting α7 nAChR are small molecules, which may interact with other nAChR subtypes and thus can suffer from selectivity issues, although some small molecules selective to α7 nAChR are also known (12). Peptide ligands pose a feasible alternative to small molecule ligands in targeting α7 nAChR due to their improved selectivity profile in targeting nAChR subtypes with potentially fewer side effects (13). Therefore, understanding the mechanism of action of peptide ligands of α7 nAChR may allow us to design better therapeutics targeting this receptor.

### α-conotoxin Mr1.7 variants target α7 nAChR

A group of natural peptides targeting nAChR are α-conotoxins. α-conotoxins are peptides found in *Conus* species consisting of 13-19 amino acids with multiple disulfide bonds that constrain their conformations (14). Several α-conotoxins have been shown to selectively interact with nAChR including α7 nAChR (15). α-conotoxins Mr1.7a (MrIA), Mr1.7b (MrIB), and Mr1.7c (MrIC) are 16-19 amino acid variants of the neuropeptide Mr1.7 (16–18). The most remarkable difference between these three peptides is their N-terminal sequence (Table 1). MrIA has a negatively-charged glutamate residue on its N-terminus, whereas MrIB has a positively-charged arginine residue. MrIC has an N-terminal proline residue, atypical for α-conotoxins. These three peptides were previously synthesized and tested for activity at α7 nAChR upon co-application with a type II positive allosteric modulator (PAM) (17, 19). Type II PAMs do not induce a response when applied by themselves, but they recover α7 nAChR from its desensitized D_s_ state, resulting in large responses to orthosteric and allosteric agonists (20, 21). Of the three peptide molecules tested, MrIA had weak partial agonist activity and MrIB was found to have no effect. MrIC (Supporting Figure 1) had no standalone activity, yet activated α7 nAChR when co-applied with the type II PAM PNU-120596 (22). However, the mechanism of activation by MrIC has not been elucidated so far.

**Table 1.** The sequences of the peptides MrIA, MrIB, and MrIC that were used previously to investigate the role of N-terminal peptide residues. The residue “O” indicates hydroxyproline.

| Peptide | Sequence |
| --- | --- |
| MrIA | ECCTHPACHVSNPELC-NH<sub>2</sub> |
| MrIB | ROECCTHOACHVSNPELCS-OH |
| MrIC | PECCTHPACHVSNPELC-NH<sub>2</sub> |

### α7 nAChR activation by MrIC may be through an allosteric binding site

Endogenous ligands of the α7 receptor bind to a site under the C-loop called “the orthosteric site”, then triggering a cascade of structural motions that result in opening of the ion channel in the transmembrane domain (TMD) (23). However, a vestibular binding site in the extracellular domain (ECD) named the “allosteric activation” (AA) site has been proposed to follow an alternative α7 nAChR activation mechanism (Figure 1). According to this hypothesis, the ligands that bind to the AA site can couple with the TMD directly and induce channel opening in the presence of a type II PAM (24–26). The MrIC activity profile is also consistent with that of allosteric agonists (17, 19). Therefore, we hypothesized that the PAM-dependent activity of MrIC may be related to the AA site of the receptor. To test this hypothesis, we first docked MrIC to the α7 nAChR along with the non-allosteric Mr1.7 variants (MrIA and MrIB) to identify the interactions that explain the unique mode of action of MrIC. Based on the identified differences, we designed three *in silico* point mutants at the AA site and docked MrIC to predict their effects on MrIC activity. Finally, MrIC activity at the three mutations was tested by voltage-clamp electrophysiology experiments to confirm the computational predictions.

**Figure 1.**
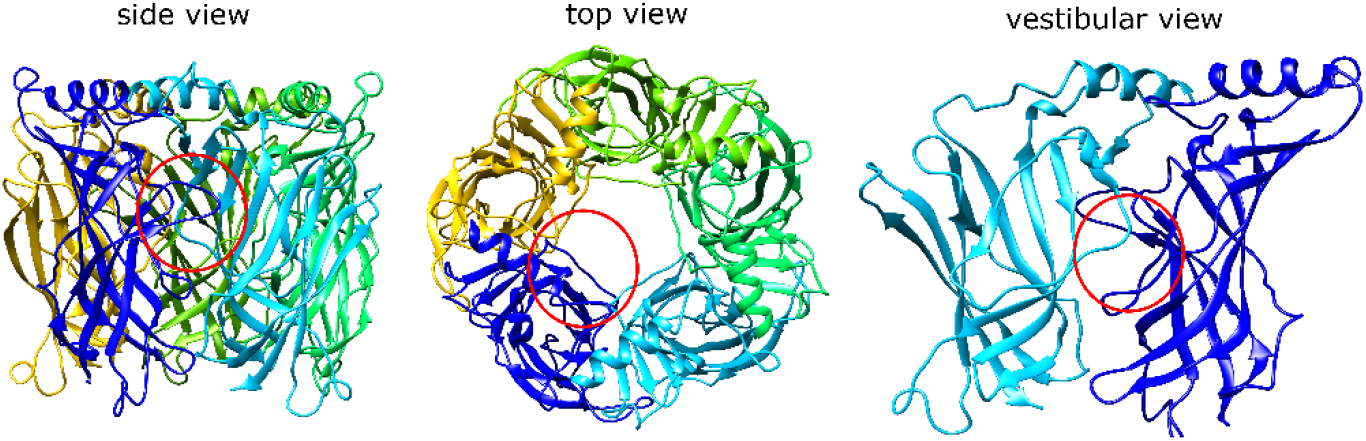
The α7 nAChR ECD structure viewed from side (left), top (middle), and the vestibule (right). Each α7 nAChR subunit is represented with a different color. The red circles indicate the position of the AA site from each perspective.

## Results and Discussion

### MrIA, MrIB, and MrIC were docked into the AA site through Rosetta peptide docking

We first docked MrIA, MrIB, and MrIC into the ECD AA site of α7 nAChR and tested whether the three molecules could bind to this site. The ECD AA site ligands investigated so far in the literature interact with this site by burying their hydrophobic parts deep into the cavity defined by the vestibular loop residues 87-93 and 98-106 (21, 25–27). The bottom of the cavity is formed by the α7 nAChR residues F33, L56, Q57, M58, I90, and L91. The charged/polar ligand residues interact with the “mouth” of the cavity, characterized by the residues R99 and D101.

Because the vestibular loops and the peptide backbones can show structural flexibility upon binding, we followed a protocol that takes such changes in flexibility into consideration. The ToxDock protocol was developed to model α-conotoxin – nAChR interactions (28). A variant of this protocol was also used to model three-finger toxin – nAChR interactions (29), showing the suitability of Rosetta peptide docking to model peptide binding to nAChR. Briefly, the peptides were first placed manually at the AA site and were subjected to a fixed-backbone relax calculation to relieve energetic frustrations. Next, the peptide – protein complexes were relaxed and a total of 200 structures were generated to sample different vestibular loop configurations. At the docking step, the three MrIC variants were docked into the ten lowest-scoring structures obtained from the relax calculations. 500 docking runs were conducted for each protein model to a total of 5000 poses for each peptide. The lowest-scoring ten structures were selected by peptide – protein interaction scores (I_sc) and were used for the analyses.

### MrIA failed to bind in the AA site but bound close to the vestibular pore

MrIA was not buried within the binding site formed by the vestibular loop, and the bulk of the molecule was oriented towards the vestibular pore (Figure 2, left panel). This resulted in a number of strong interactions with the subunit C in addition to the subunits A and B. The strongest MrIA interaction was an electrostatic interaction between the N-terminal E1^MrIA^ and α7R99 (Supporting Figure 2A). Other interactions between MrIA residues and the α7 nAChR involved α7 nAChR H105, R20, Q84, T106, D101, Y15, A102, and G83. The interactions with α7 Q84, H105, and T106 of the subunit C were particularly strong.

**Figure 2.**
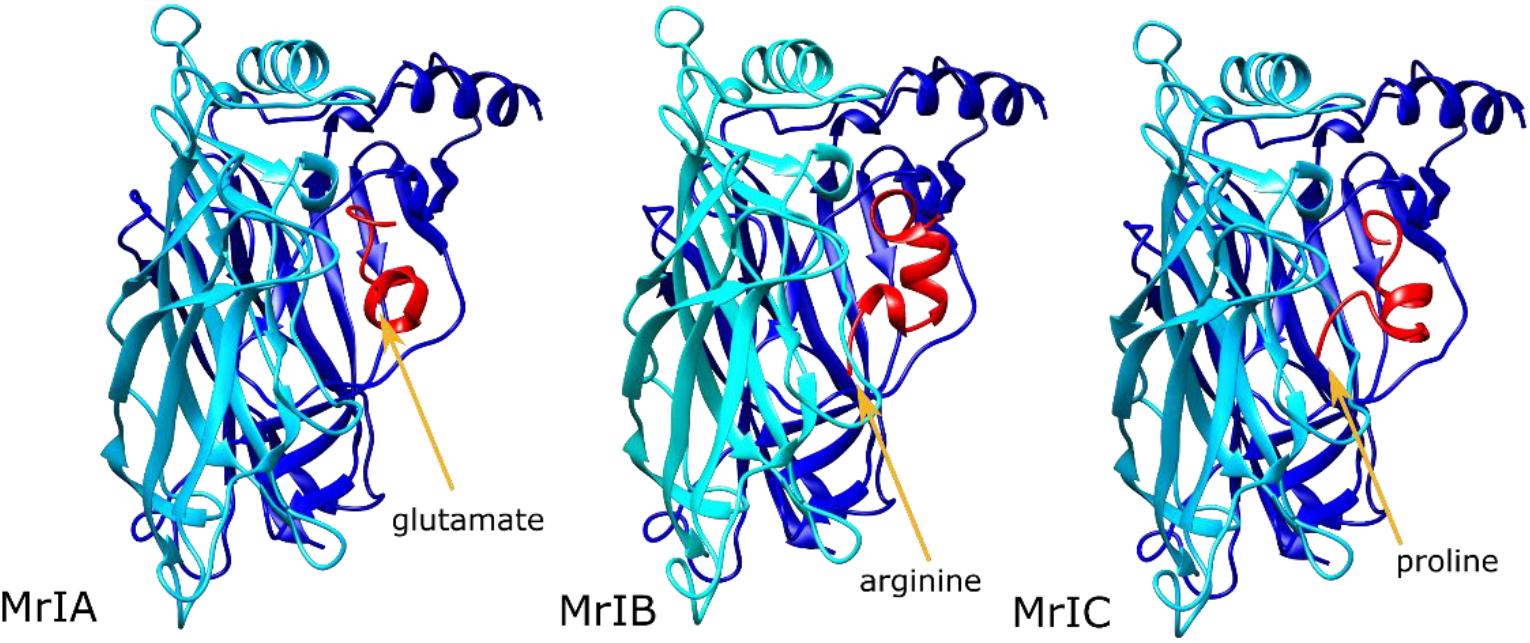
The lowest-scoring MrIA (left), MrIB (middle), and MrIC (right) poses at the α7 nAChR ECD AA site. The α7 nAChR subunits are shown in blue (Subunit A) and cyan (Subunit B), and the peptides are shown in red. The orange arrows indicate the positions of the N-terminal residue of each peptide.

### MrIB shifted the vestibular loops away from each other

In the MrIB calculations, the N-terminal arginine residue bound between the vestibular loops of the primary and complementary interfaces, prying the vestibular loop of the negative interface away from that of the positive interface (Figure 2, middle panel). This resulted in MrIB interacting with the AA site of a single subunit, but it kept contact with the vestibular loop residues from both subunits. Favorable MrIB interactions were formed with the α7 residues D101, A102, G83, T103, T106, Q84, and F100. The two strongest interactions were hydrophobic interactions between A9^MrIB^ and α7A102, and a hydrogen bond between H7^MrIB^ and α7D101. H7^MrIB^ also formed a hydrogen bond with the α7R99 residue, which interacted with the peptide residue E1^MrIA^ in the MrIA calculations. Contrary to MrIA, α7R99 interactions were present only at a single subunit (Supporting Figure 2B).

### MrIC anchored the AA site through its N-terminal proline

MrIC interacted with the AA sites of two interfaces differently from MrIA and MrIB. One of these AA site interactions was formed through binding of the α-helical portion of the peptide (residues 7-14) to the cleft between the subunits A and B. The other interaction with the AA site was formed between the subunits B and C through anchoring of P1^MrIC^ residue at this site, although there were no significant interactions with the C subunit side, unlike MrIA (Figure 2, right panel). Binding at the second AA site resulted in pulling the vestibular loop of the subunit B towards the vestibular loop of the subunit A, therefore strengthening the interactions between the two loops. P1^MrIC^ bound among the α7 residues P121, S95, and D97, but formed only hydrophobic interactions these residues.

The α7 nAChR residues that had an interaction energy with the peptide better than −1 Rosetta Energy Units (REU) were R99, Y15, D101, F104, A102, G83, P81, H105, and T106. Of these residues, R99 had the lowest interaction score, owing to a strong electrostatic interaction between this residue and E2^MrIC^. The second strongest interaction was a hydrogen bond between α7Y15 and E15^MrIC^. α7D101 formed a hydrogen bond with H6^MrIC^ at subunit A and a backbone-backbone hydrogen bond with E2^MrIC^ at the subunit B. These observations suggest that the role of P1^MrIC^ may be placement of E2^MrlC^ close to α7R99 in a stable manner rather than forming interactions within the AA site by itself (Supporting Figure 2C).

Overall, our results demonstrated that the N-terminal proline residue of MrIC is a suitable anchor point for the AA site, placing E2^MrIC^ in close contact with α7R99. Contrarily, the E1^MrIA^ residue could not anchor to the AA site, and R1^MrIB^ clashed with α7R99 at the mouth of the vestibular site. Lack of MrIA and MrIB binding at the AA site is consistent with the non-allosteric modes of action observed for the two peptides.

### MrIC binding to the orthosteric site was insignificant in electrophysiology experiments

We started our experimental investigation by testing the binding of MrIC to the α7 nAChR orthosteric site in experiments with the wild-type (WT) α7 nAChR. First, we co-applied 60 μM acetylcholine (ACh) and 50 μM MrIC to test whether MrIC could compete with ACh at the orthosteric site. The results showed effectively no change in ACh responses at this MrIC concentration (Supporting Figure 3A). Second, we co-applied 60 μM PNU-120596 + 50 μM MrIC in the absence and presence of 100 μM (−)2,3,5,6TMP-TQS ((−)TMP-TQS, Supporting Figure 1), a selective antagonist of the ECD AA site that has little effect on the orthosteric and TMD PAM site activity of α7 nAChR (27, 30).

Standalone applications of 50 μM MrIC or 60 μM PNU-120596 (data not shown) applied to WT α7 nAChR evoked no response, consistent with the previous observations in SH-S5Y5 cells (17, 19). On the other hand, co-application of 50 μM MrIC with 60 μM PNU-120596 evoked a net-charge response 2.13 times the ACh controls. Application of (−)TMP-TQS caused ∼90% diminution of the MrIC-potentiated response, which was statistically significant (Supporting Figure 3B). The lack of ACh competition and (−)TMP-TQS-dependent loss of potentiated MrIC activity suggests lack of orthosteric binding under these conditions.

### Upon co-application with PNU-120596, MrIC could activate the α7C190 mutant, which cannot be orthosterically activated

Based on the results with the WT receptor, we tested MrIC activity at the α7C190 mutant. α7 nAChR mutants such as C190A are called “non orthosterically activatible receptors” (NOARs) because they cannot be activated by traditional agonists of the α7 nAChR, including ACh. Although NOARs cannot be activated orthosterically, they can be activated by standalone applications of agonist-PAMs (ago-PAMs) or allosteric agonists plus type II PAMs (21, 25, 31). Because α7C190A can only be activated allosterically, we tested MrIC responses with this mutant to provide further support for the allosteric activation hypothesis for MrIC. Since it is not possible to normalize the MrIC responses with ACh applications to this mutant, we used the ago-PAM GAT107 (Supporting Figure 1)(32, 33) activity as the reference to normalize the MrIC responses.

As expected, standalone application of 50 μM MrIC had no effect on α7C190A (data not shown). 1 μM GAT107 gave a large response, which was inhibited by the application of 100 μM (−)TMP-TQS (Figure 3, left panel). Co-application of 60 μM PNU + 50 μM MrIC evoked a response that was also diminished by the application of 100 μM (−)TMP-TQS (Figure 3, right panel). In summary, the WT and α7C190A activity profiles of MrIC confirmed that MrIC acts as an allosteric agonist of α7 nAChR. We then sought to confirm that MrIC acts through the ECD AA site by testing for changes in MrIC activity caused by the AA site mutations.

**Figure 3.**
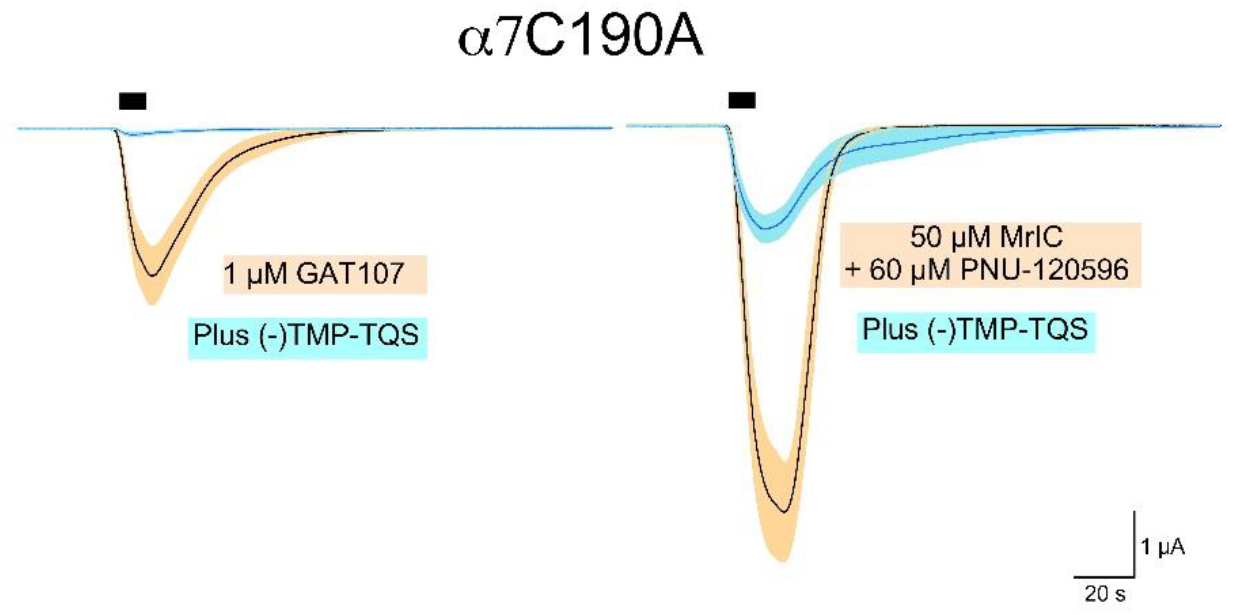
Left panel: The α7C190A activity induced by 1μM GAT107 in the absence (black line) and presence (blue line) of (−)100 μM TMP-TQS. Right panel: The activity induced by 60 μM PNU + 50 μM MrIC in the absence (black line) and presence (blue line) of 100 μM (−)TMP-TQS. The black box indicates the time of MrIC application. The orange and cyan bands indicate the error margins of the measurements.

### MrIC binding to the ECD AA site was tested with three AA site mutants

Further support for MrIC binding at the AA site was provided by mutagenesis experiments. AA site mutations typically have little or no effect on orthosteric activation, but result in large variations in allosteric activation. Mutations can be designed to increase (gain-of-activity) or decrease (loss-of-activity) the allosteric responses of α7 nAChR. AA site mutations such as D101A are known to diminish allosteric activation (25). However, loss-of-activity may be caused by a variety of factors independent of the bound ligand. As a result, mutations that cause an increase in activity provide stronger support for the involvement of the AA site.

We focused on identifying mutants that could improve binding of MrIC. Computational studies to improve α-conotoxin binding to the α7 orthosteric site demonstrated the importance of hydrophobic interactions between α-conotoxins and the α7 binding site (34). Based on this logic, α7G83V and α7A102V were predicted to increase MrIC activity since they replace a small residue with a larger hydrophobic residue, creating additional contact surface for interactions between the peptide and the protein. The third mutation selected was α7T106A. The α7T106 residue forms hydrophobic interactions with V11^MrIC^ and L16^MrIC^ and blocks the complete entry of the peptide into the AA site formed between the α7 nAChR subunits A and B. This mutation is also of interest since it responds to standalone applications of PNU-120596 in a (−)TMP-TQS-sensitive manner, suggestive of a role in receptor desensitization (27). Therefore, MrIC was docked into the three mutant receptors to investigate the different interactions associated with these mutations. The locations of these residues and their vicinities can be seen in Supporting Figure 4.

### The α7G83V mutation caused an increase in the binding strength of several peptide residues in peptide docking calculations

The α7G83V mutation resulted in changes in identity and strength of the peptide interactions with the protein. Because of the increased bulk of the binding site at a region close to the C-terminus of the peptide, the peptide C-terminus extended away from the AA site, pointing toward the vestibular space (Figure 4A). The average interaction score calculated for the α7G83V poses was 2.8 REU lower than the WT poses (Supporting Table 1). Instead of a large increase caused by the lower scores of a few residues, there was a small but consistent decrease in the scores of several residues. The strongest interaction was an electrostatic interaction between α7R99 and E2^MrIC^ as observed for the WT calculations. The backbone of P1^MrIC^ formed a hydrogen bond with the -OH group of α7T103, different than the WT calculations. α7D101 showed backbone-backbone hydrogen bonding with E2^MrIC^. The H6^MrIC^ side chain formed a hydrogen bond with the α7T103 backbone in some poses and interacted with α7D101 in others. α7K87 moved closer to the peptide and formed a hydrogen bond with the backbone of A8^MrIC^. α7V83 formed weak hydrophobic interactions with P14^MrIC^, and interacted with the side chain of E15^MrIC^. E15^MrIC^ also formed a hydrogen bond with α7Q84.

**Figure 4.**
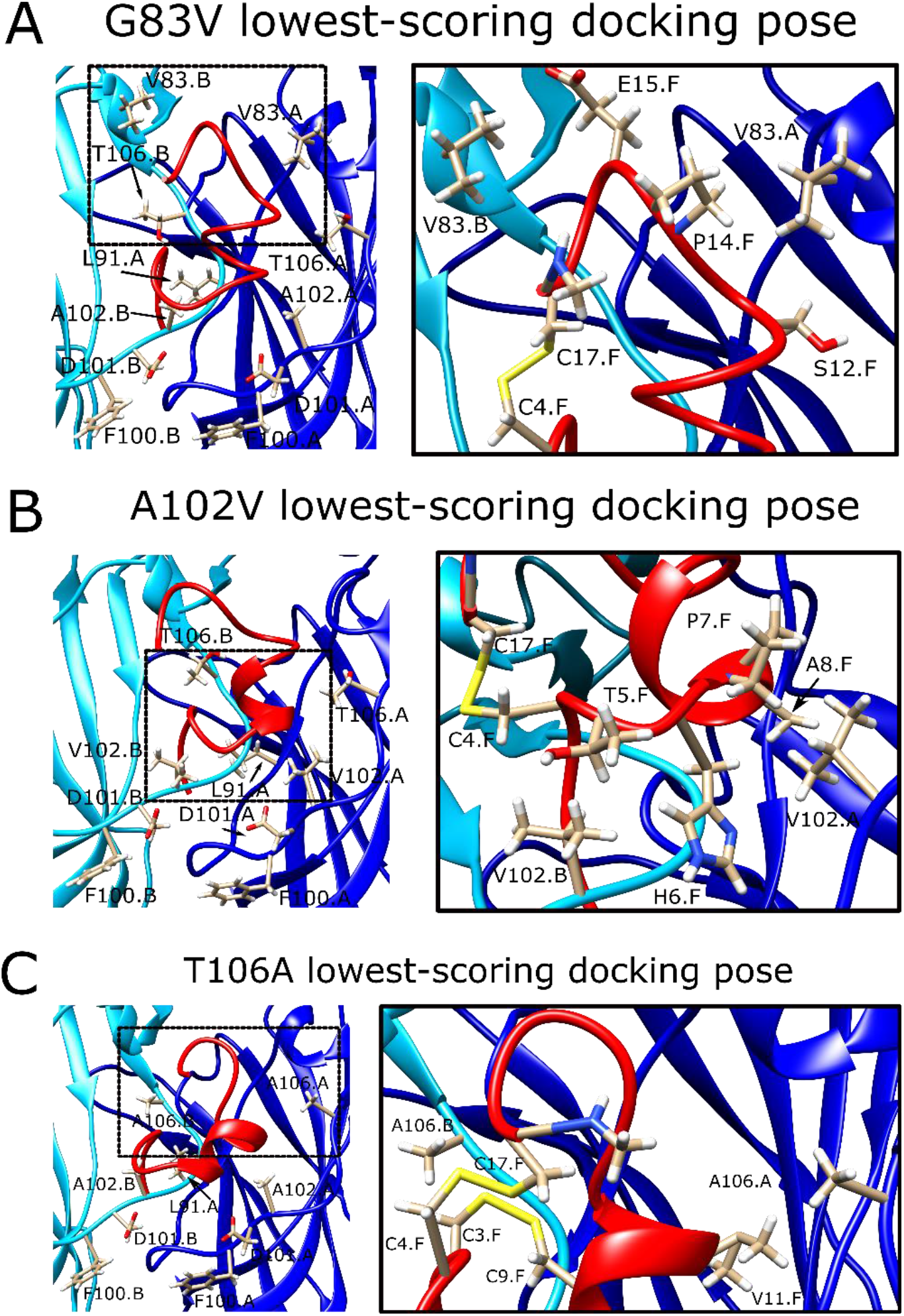
The lowest-scoring docking poses of MrIC at the α7G83V (A), α7A102V (B), and α7T106A (C) vestibular sites viewed from the vestibule with important residues labeled (left panels), and close-ups of the dashed areas around the mutated residues, showing specific interactions between MrIC and the vestibular residues (right panels). Chain A (blue) stands for the positive subunit of the interface bearing the C-loop, chain B (cyan) stands for the negative subunit of the interface, and chain F (red) stands for the peptide.

### The α7A102V mutation improved the binding strength slightly less than the α7G83V mutation

The α7A102V mutation affected both the conformation of MrIC and the interactions between the peptide and the protein. In order to adapt to the bulk introduced by α7V102 at the subunit B, MrIC bent away from this residue at the peptide residues C4^MrIC^ and T5^MrIC^, which replaced the N-terminal residues of the peptide, although P1^MrIC^ remained anchored at the vestibular site (Figure 4B). The strongest interaction was between α7R99 of the subunit B and E2^MrIC^, similar to the calculations for WT α7 nAChR. The α7A102V mutation caused loss of interactions at a single peptide residue (V11^MrIC^), but resulted in the formation of four additional interactions between the peptide and α7V102 (C4^MrIC^, H6^MrIC^, P7^MrIC^, A8^MrIC^). Of these residues, H6^MrIC^ also formed hydrogen bonds with α7E98 and α7R99. In the absence of α7A102, V11^MrIC^ formed new interactions with the side chains of α7D82 and α7T106. The average interface score of the top ten poses was 2.1 REU lower than the WT α7 nAChR score (Supporting Table 1).

### α7T106A peptide docking poses were similar to the WT α7 nAChR with only minor improvement of interaction energy

The decrease observed for the average interface score of the α7T106A calculations was modest compared to the α7G83V and α7A102V calculations, with an improvement of only 0.9 REU (Supporting Table 1). The peptide configuration was also similar to the WT with a minor disruption in the helical geometry of its α-helical region (Figure 4C). The newly-introduced α7A106 mostly mimicked the -CH_3_ group of α7T106, but it created extra space for the peptide to bind because of its reduced size compared to T106. The α7R99 – E2^MrIC^ interaction was conserved. The α7D101 residues on both interfacing subunits interacted with the peptide residues in a way similar to the G83V mutant. Other differences compared to α7G83V and α7A102V were a more intimate interaction within the AA site characterized by the interaction between V11^MrIC^ and α7P121. α7H105 formed a hydrogen bond with the -OH group of S12^MrIC^.

### α7G83V and α7A102V mutations caused increased MrIC responses compared to WT α7 nAChR in electrophysiology experiments

The α7G83V and α7A102V activities were tested the same way as the WT α7 nAChR experiments. Both mutations showed negligible responses to standalone 60 μM PNU-120596 applications. Co-application of 50 μM MrIC and 60 μM PNU-120596 evoked 26-fold and 94-fold increase in net charge compared to the WT α7 nAChR MrIC responses for α7G83V and α7A102V, respectively (Figure 5, left and middle panels). Based on these results, increasing the hydrophobic surface of the AA site causes an increase in potentiated responses.

**Figure 5.**
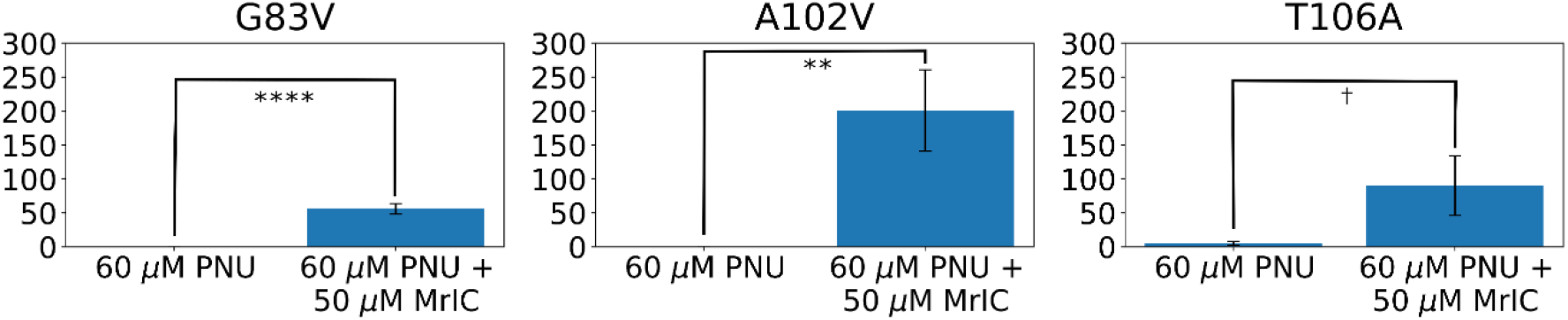
The α7G83V (left), α7A102V (middle), and α7T106A (right) responses evoked by the application of 60 μM PNU-120596 or 50 μM MrIC + 60 μM PNU-120596. All responses were normalized to applications of 60 μM ACh. † stands for p < 0.10, ^*^ stands for p < 0.05, ^* *^stands for p < 0.01, ^***^stands for p < 0.001, and ^****^ stands for p < 0.0001.

### The α7T106A mutation resulted in background responses to PNU-120596 with an enhancement caused by MrIC

The α7T106A mutation resulted in an increased background net charge response to 60 μM PNU-120596 (∼6-fold), contrary to the results with the WT α7 nAChR, α7G83V, and α7A102V receptors (Figure 5, right panel). This suggests that the changes to the vestibular loop conformation induced by the α7T106A mutation may put α7 nAChR into a desensitized state independent of ligand binding. The activity increase caused by MrIC was ∼15 times compared to the α7T106A PNU-120596 application and 42 times the WT α7 nAChR response to MrIC + PNU-120596 application.

In summary, all three predicted AA site mutations resulted in increased MrIC activity, consistent with allosteric activation through this site. These results combined with MrIC’s lack of ACh competition, (−)TMP-TQS-sensitivity, and the ability to activate the NOAR α7C190A strongly suggest that the source of MrIC allosteric activity is the vestibular AA site. The unique allosteric behavior of MrIC among other Mr1.7 variants may be connected to its N-terminal proline, which can anchor to the AA site and induce electrostatic interactions between E2^MrIC^ and α7R99 residues. Our study sheds light on how α-conotoxins can activate the α7 nAChR through allosteric mechanisms and underline the value of computational methods in predicting the mode of action of peptide ligands. Further, the mechanistic insights provided by our study pave the way for the design of peptide drugs for allosteric activation of α7 nAChR.

### Limitations of the study

Changes in the conformations of the α7 nAChR AA site and the α-conotoxins prevent the determination of a single conformation that represents the interaction between the peptide and the receptor. In order to address this issue, we averaged energies from the ten lowest-scoring poses to calculate the peptide binding energies. Although this approach was useful to predict mutations that increase MrIC binding, it was unable to make quantitative predictions regarding activity increase since the predicted order of activities was different than the experimentally-determined values. More comprehensive studies to determine MrIC activity at a range of concentrations may be useful to assess the predictive utility of the docking methodology used in this study.

Our findings are different than the previous results whereby MrIC only acted as an antagonist at *Xenopus* oocytes with no response to PNU-120596 co-application (17, 19). This discrepancy may be caused by differences of the applied MrIC and PNU-120596 concentrations, isomerization of MrIC in solution, or differences in peptide preparation that may have resulted in a mixture of active and inactive isomers. Therefore, extended analyses on isomerization of MrIC in solution and the activity of different MrIC isomers are required to understand the variations observed in different experimental setups.

## Materials and Methods

### Homology modeling of α7 ECD

The α-conotoxin ImI-bound (PDB ID: 2C9T) (35), and lobeline (orthosteric) and fragment 5-bound (AA site) (PDB ID: 5AFN) (36) acetylcholine binding protein (AChBP) structures were used as the templates for the α7 ECD homology model used for the docking calculations with a query of mature α7 ECD sequence (Uniprot ID: P36544, CHRNA7). 3- and 9-mer fragments were generated locally, and the PSIPRED server was used for the prediction of the secondary structure of the target sequence. All disulfide bonds were identified based on the bonding patterns of the template structures. A total of 1000 structures were generated using the *ref2015_cart* score function (37, 38) and fivefold symmetry was imposed on the models. The lowest-scoring structure was selected as the docking model. An apo (no ligand bound) homology model was also created as a reference point for the rotamer configurations in the absence of a peptide. This structure was used for the peptide docking step as the unbound model. The same protocol was followed to generate the apo structure with the difference that the template used for this structure was the apo α7-AChBP structure (PDB ID: 3SQ9, (39)).

### Modeling of MrIA, MrIB, and MrIC

All three structures were modeled based on the structure of α-conotoxin OmIA (PDB ID: 2GCZ) (40). The mutations to obtain the corresponding target peptide sequences (**Table 1**) were introduced with UCSF Chimera (41). For MrIB, the hydroxyproline residues were replaced with proline residues due to the lack of parameters for this non-canonical amino acid. Each peptide structure was relaxed prior to the docking calculations using the *talaris2013* score function (42). The relaxed structures were used for the docking calculations.

### Docking of the peptides into α7 nAChR models

A modified version of the ToxDock protocol (28) was used for the peptide docking calculations. Specifically, the peptide – α7 ECD complexes were manually generated for MrIA, MrIB, and MrIC. The MrIA complex was generated first and the remaining structures were aligned on this structure to ensure a consistent starting point for all three peptides. For the G83V, A102V, and T106A mutations, UCSF Chimera was used to introduce the point mutations at the relevant subunits through the “Rotamers” menu. A fixed-backbone relax calculation was run to relieve any energetic frustrations, followed by Rosetta relax calculations (43) to generate 200 models. The top 10 structures were selected as the starting point for the docking calculations based on the lowest total scores. FlexPepDock (44) was used to dock the peptides into the AA site. 500 docking poses were generated for each starting point, resulting in a total of 5000 structures for each peptide. The lowest-scoring ten poses were selected based on lowest interface (I_sc) scores and used for visual inspection and score comparison. All calculations were run with the *talaris2013* score function.

### Chemicals and reagents

Acetylcholine chloride (ACh) and buffer chemicals were purchased from Sigma-Aldrich Chemical Company (St. Louis, MO). PNU-120596 was synthesized in Dr. Nicole A. Horenstein’s laboratory (University of Florida, Gainesville, FL, USA) by Dr. Kinga Chojnacka following the published procedure by Hurst et al. 2005 (22). GAT107 and (−)TMP-TQS were provided by Dr. Ganesh Thakur (Northeastern University, Boston, MA, USA) following the procedure by Thakur et al. 2013 and Kulkarni et al. 2013 (33, 45). While preliminary studies and the identification of the active isomer were conducted with toxin synthesized by the Vetter laboratory as described in the following section, all of the data in the figures were obtained with MrIC purchased from Alomone Laboratories (Jerusalem, Israel, lot number STC320TX011).

### Peptide regioselective synthesis and purification

MrIC was synthesized by standard Fmoc chemistry with Cys side-chains protected as one pair ACM (1,3 or 1,4) and the other pair Trt (2,4 or 2,3), starting from Polystyrene AM RAM resin on a 0.62 mmol scale. The deprotection of the side chain group and cleavage of the linear peptide from the resin were conducted with a TFA (92.5%): H_2_O (2.5%): TIPSi (2.5%): DODt (2.5%) solution for 40 mins at 40 °C. Linear peptide was precipitated and washed with cold ether, dissolved in 45% acetonitrile/0.05% TFA/H_2_O, filtered and lyophilized on a freeze dryer. The crude linear peptide was purified by preparative RP-HPLC (Vydac C18). For the one-pot regioselective formation of the 1-3 and 2-4 (or 1-4, 2-3) disulfide bonds, the method has been described before (46). Briefly, the crude dithiol product was first subjected to a 5 min I_2_ treatment in 90% AcOH / MeOH to form the Cys 2-4 or Cys 2-3 disulfide bond. An aliquot was analysed by HPLC and HRMS to confirm the complete formation of the first pair disulfide bond and the integrity of the Cys 1,3 or Cys 1,4 Acm groups. To induce the removal of the Acm groups and form the Cys 1-3 or Cys 1-4 disulfide bond, 1% TFA / H_2_O was added followed by additional I_2_ and the reaction was allowed to proceed for a further 90 min, giving Globular Cys1-3, 2-4 MrIC. Then, the oxidized peptide was purified by HPLC.

### Heterologous expression of nAChRs in *Xenopus* laevis oocytes

The human α7 nAChR clone was obtained from Dr. J. Lindstrom (University of Pennsylvania, Philadelphia, PA). The human resistance-to-cholinesterase 3 (RIC-3) clone was obtained from Dr. M. Treinin (Hebrew University, Jerusalem, Israel) and co-injected with α7 to improve the level and speed of α7 receptor expression without affecting the pharmacological properties of the receptors (47). Subsequent to linearization and purification of the plasmid cDNAs, cRNAs were prepared using the mMessage mMachine in vitro RNA transcription kit (Ambion, Austin, TX). The α7C190A mutant was made as previously described with a C116S double mutation to prevent spurious disulfide bond formation with the free cysteine (48).

Oocytes were surgically removed from mature female Xenopus laevis frogs (Nasco, Ft. Atkinson, WI). Frogs were maintained in the Animal Care Service facility of the University of Florida, and all procedures were approved by the University of Florida Institutional Animal Care and Use Committee. In brief, the frog was first anesthetized for 15-20 min in 1.5 liters frog tank water containing 1 g of 3-aminobenzoate methanesulfonate (MS-222) buffered with sodium bicarbonate. The harvested oocytes were treated with 1.4 mg/ml Type 1 collagenase (Worthington Biochemicals, Freehold NJ) for 2-4 h at room temperature in calcium-free Barth’s solution (88 mM NaCl, 1 mM KCl, 2.38 mM NaHCO_3_, 0.82 mM MgSO_4_, 15 mM HEPES, and 12 mg/l tetracycline, pH 7.6) to remove the ovarian tissue and the follicular layers. Stage V oocytes were subsequently isolated and injected with 50 nl of 5-20 ng nAChR subunit cRNA. Oocytes were maintained in Barth’s solution with calcium (additional 0.32 mM Ca(NO_3_)_2_ and 0.41 mM CaCl_2_), and recordings were carried out 2-14 days after injection.

### Two-electrode voltage clamp electrophysiology

Experiments were conducted using OpusXpress 6000A (Molecular Devices, Union City, CA) (49). Both the voltage and current electrodes were filled with 3 M KCl. Oocytes were voltage-clamped at −60 mV at room temperature (24°C). The oocytes were bath-perfused with Ringer’s solution (115 mM NaCl, 2.5 mM KCl, 1.8 mM CaCl_2_, 10 mM HEPES, and 1 μM atropine, pH 7.2) at 2 ml/min. To evaluate the effects of experimental compounds compared to ACh-evoked responses of α7 nAChR subtypes expressed in oocytes, control responses were defined as the average of two initial applications of ACh made before test applications. Solutions were applied from 96-well plates via disposable tips. Drug applications were 12 s in duration followed by 181 s washout periods. A typical recording for each set of oocytes constituted two initial control applications of ACh, one or more experimental compound applications, and then a follow-up control application(s) of ACh. The control ACh concentration was 60 μM for the wild type (WT) and the three AA site mutant receptor experiments, and the average of independent 1 μM GAT107 responses were used as the control for the NOAR experiments. The responses were calculated as both peak current amplitudes and net charge, as previously described (50), and the averages of the two initial controls were used for normalization purposes. Data were collected at 50 Hz, filtered at 20 Hz, and analyzed by Clampfit 9.2 or 10.0 (Molecular Devices) and Excel (Microsoft, Redmond, WA). Data were expressed as means ± SEM from at least four oocytes for each experiment, unless otherwise stated, and plotted by matplotlib.pyplot library of Python. Multi-cell averages were calculated for comparisons of complex responses. Averages of the normalized data were calculated for each of the 10,322 points in each of the 206.44 s traces (acquired at 50 Hz), as well as the standard errors for those averages.

### Data and statistical analysis

Comparisons of results were made using t-tests between the pairs of experimental measurements. A value of p <0.10 was used to constitute the level of significance. The statistics were calculated using an Excel template provided in Microsoft Office. For some experiments statistical comparisons were not calculated but the data plotted as the point-by-point averages (± SEM), permitting qualitative evaluation of the data by visual inspection.

## Acknowledgments

We thank Dr. Nicole A. Horenstein for providing PNU-120596 and her comments on the manuscript. We thank Dr. Ganesh Thakur for providing GAT107 and (−)TMP-TQS.

## Figures and Tables

**Supporting Table 1.** The average scores and standard deviations (S.D.) calculated for the ten lowest-scoring poses from the WT, G83V, A102V, and T106A peptide docking calculations. All units are in Rosetta Energy Units (REU).

| Protein | Average Score (REU) | S.D. (REU) |
| --- | --- | --- |
| WT | -25.9 | 0.8 |
| G83V | -28.7 | 0.8 |
| A102V | -28.0 | 0.6 |
| T106A | -26.8 | 1.0 |

**Supporting Figure 1.**
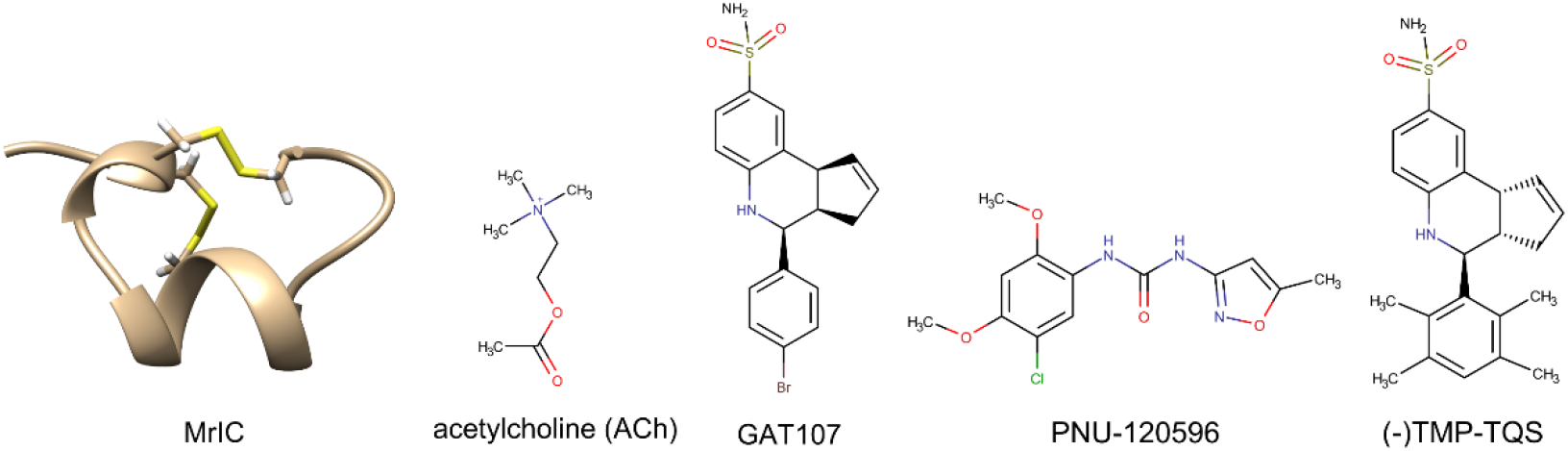
The structures of an MrIC model, ACh, GAT107, PNU-120596, and (−)TMP-TQS that were used in the experimental and computational studies.

**Supporting Figure 2.**
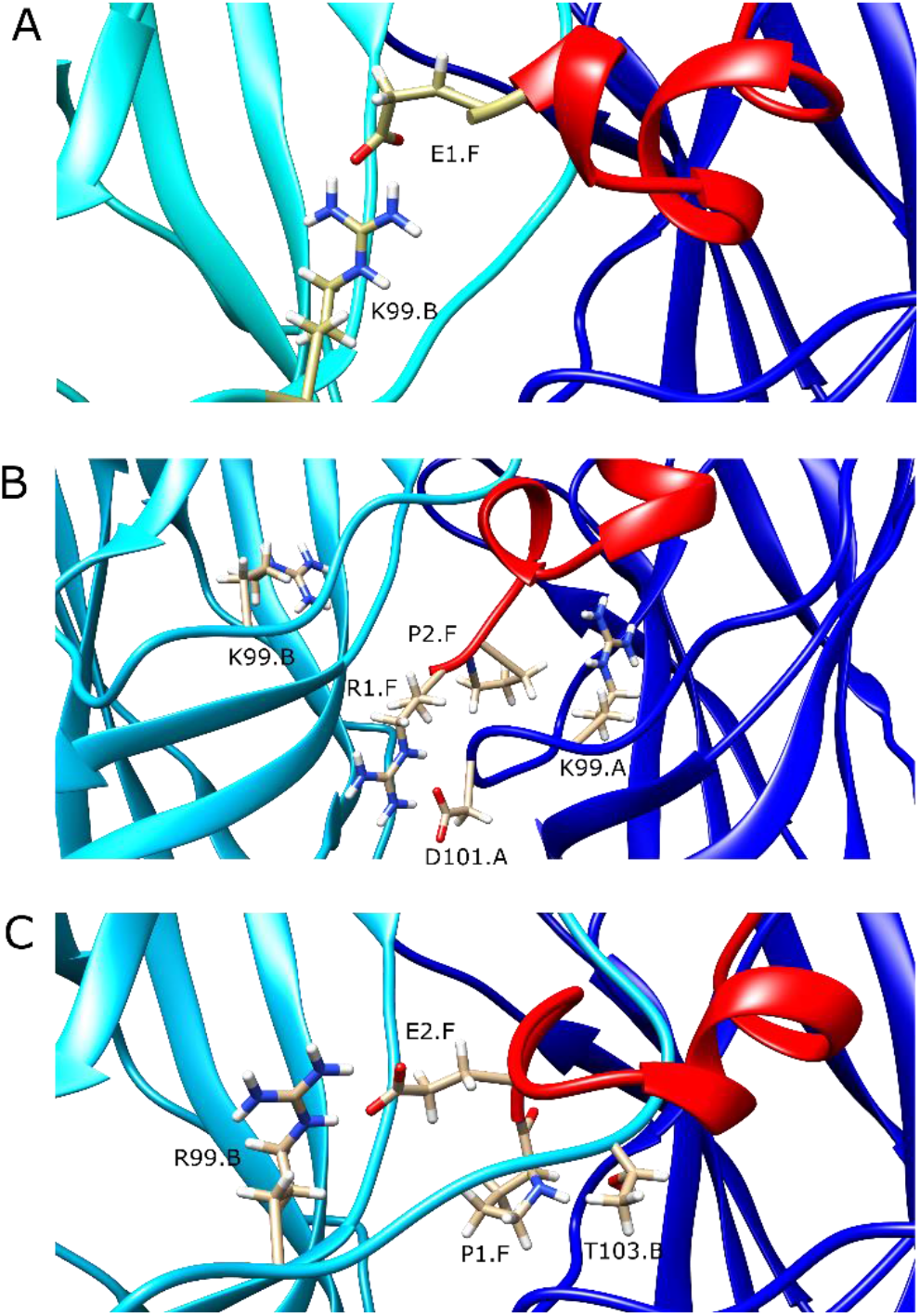
The interactions between MrIA (A), MrIB (B), and MrIC (C) and the vestibular residues. Chain A (blue) indicates subunit A, chain B (cyan) indicates subunit B, and chain F (red) indicates MrIC. The residues important for α7 nAChR – MrIC interactions are shown explicitly.

**Supporting Figure 3.**
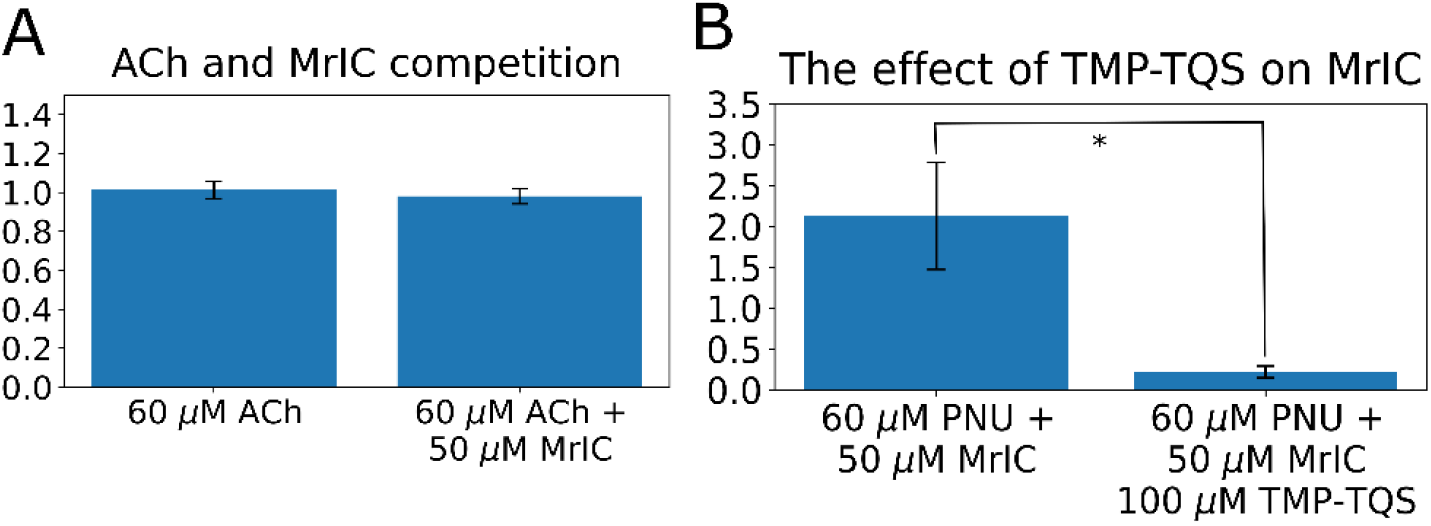
The effect of 50 μM MrIC application on the activity of 60 μM ACh (A), and the effect of 100 μM (−)TMP-TQS on the potentiated response evoked by 60 μM PNU-120596 + 50 μM MrIC. All responses were normalized to the responses evoked by 60 μM ACh applications. † stands for p < 0.10, *stands for p < 0.05, **stands for p < 0.01, ***stands for p < 0.001, and ****stands for p < 0.0001.

**Supporting Figure 4.**
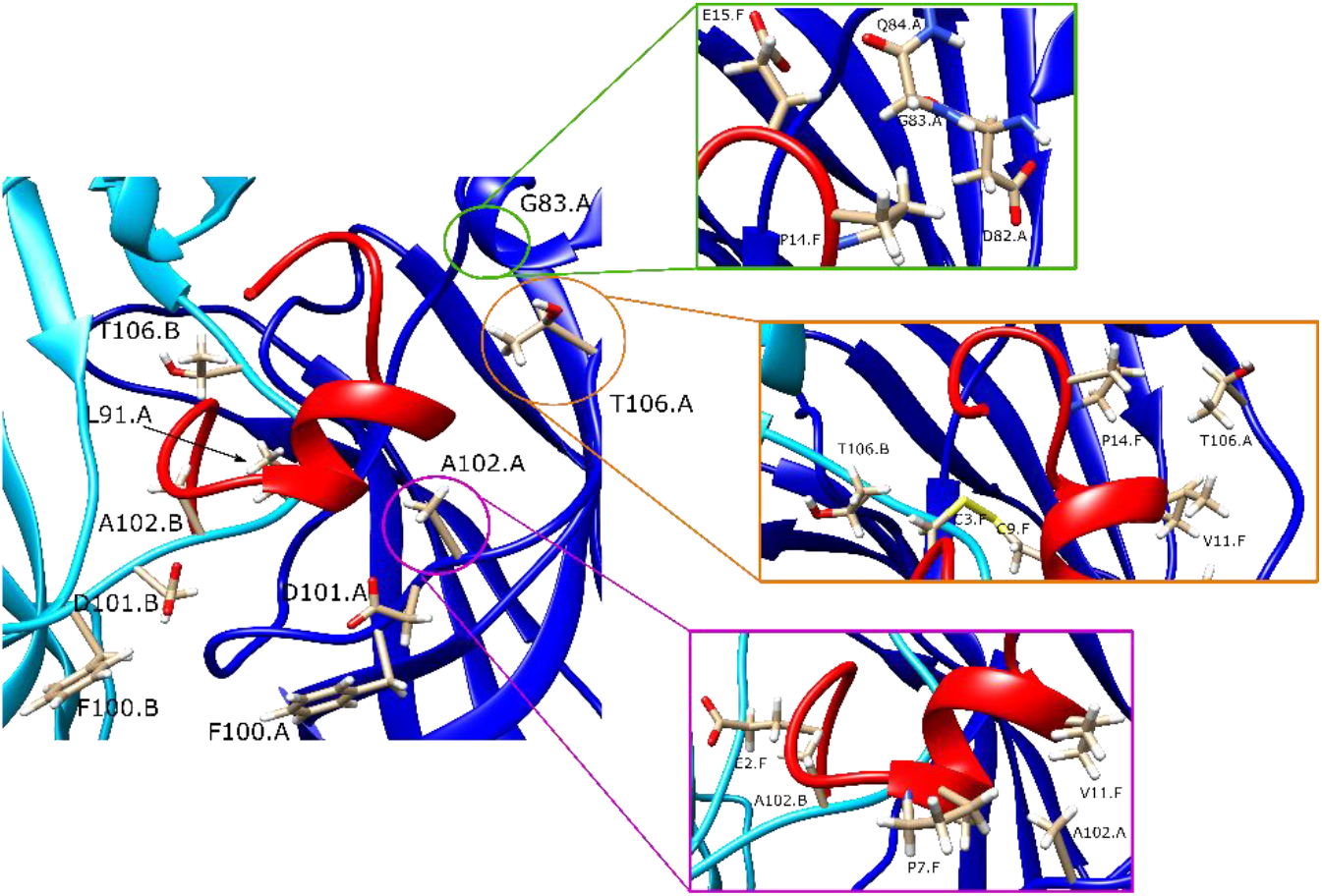
The lowest-scoring MrIC pose at the WT α7 nAChR AA site and the peptide-protein interactions at the vicinity of the residues G83 (green box), T106 (orange box), and A102 (purple box). The chain A (blue) indicates subunit A, the chain B (cyan) indicates subunit B, and the chain F (red) indicates MrIC. The residues important for α7 nAChR – MrIC interactions are shown explicitly.

